# Neutrophil-specific STAT4 deficiency attenuates atherosclerotic burden and improves plaque stability via reduction in neutrophil activation and recruitment into aortas of *Ldlr*^-/-^ mice

**DOI:** 10.1101/2023.02.22.529608

**Authors:** W. Coles Keeter, Alina K Moriarty, Rachel Akers, Shelby Ma, Marion Mussbacher, Jerry L. Nadler, Elena V. Galkina

## Abstract

**Background and Aims:** Neutrophils drive atheroprogression and directly contribute to plaque instability. We recently identified signal transducer and activator of transcription 4 (STAT4) as a critical component for bacterial host defense in neutrophils. The STAT4-dependent functions of neutrophils in atherogenesis are unknown. Therefore, we investigated a contributory role of STAT4 in neutrophils during advanced atherosclerosis.

**Methods:** We generated myeloid-specific *Stat4*^*ΔLysM*^*Ldlr*^*-/-*^, neutrophil-specific *Stat4*^*ΔS100A8*^ *Ldlr*^*-/-*^, and control *Stat4*^*fl/fl*^*Ldlr*^*-/-*^ mice. All groups were fed a high-fat/cholesterol diet (HFD-C) for 28 weeks to establish advanced atherosclerosis. Aortic root plaque burden and stability were assessed histologically by Movat Pentachrome staining. Nanostring gene expression analysis was performed on isolated blood neutrophils. Flow cytometry was utilized to analyze hematopoiesis and blood neutrophil activation. *In vivo* homing of neutrophils to atherosclerotic plaques was performed by adoptively transferring prelabeled *Stat4*^*ΔLysM*^*Ldlr*^*-/-*^ and *Stat4*^*fl/fl*^*Ldlr*^*-/-*^ bone marrow cells into aged atherosclerotic *Apoe*^*-/-*^ mice and detected by flow cytometry.

**Results:** STAT4 deficiency in both myeloid-specific and neutrophil-specific mice provided similar reductions in aortic root plaque burden and improvements in plaque stability via reduction in necrotic core size, improved fibrous cap area, and increased vascular smooth muscle cell content within the fibrous cap. Myeloid-specific STAT4 deficiency resulted in decreased circulating neutrophils via reduced production of granulocyte-monocyte progenitors in the bone marrow. Neutrophil activation was dampened in *Stat4*^*ΔLysM*^*Ldlr*^*-/-*^ mice via reduced mitochondrial superoxide production, attenuated surface expression of degranulation marker CD63, and reduced frequency of neutrophil-platelet aggregates. Myeloid-specific STAT4 deficiency diminished expression of chemokine receptors CCR1 and CCR2 and impaired *in vivo* neutrophil trafficking to atherosclerotic aorta.

**Conclusions:** Our work indicates a pro-atherogenic role for STAT4-dependent neutrophil activation and how it contributes to multiple factors of plaque instability during advanced atherosclerosis in mice.

## Introduction

Atherosclerosis is the etiological culprit of cardiovascular disease (CVD), whose complications via myocardial infarction and stroke remain the leading cause of mortality in the developed world.^1^ Despite significant improvements in lipid lowering therapies, the clinical burden of patients with CVD is a continuously unmet challenge. Research efforts over the past two decades have revealed both the innate and adaptive arms of the immune system as key contributors to the initiation and progression of atherosclerosis.^2^ Although certain subsets of leukocytes have atheroprotective properties, the dysregulation of activated leukocytes has become a key target for novel therapeutic interventions. Neutrophils are the most abundant immune cells in the blood, accounting for approximately 40-70% of all circulating leukocytes in humans. However, neutrophils represent a minor proportion of intraplaque immune cells, which are mostly dominated by macrophages and T cells.^3^ Historically, neutrophils and their diverse armament of proinflammatory functions have remained largely ignored in the study of atherosclerosis. However, recent studies have demonstrated targeting neutrophilia and inhibiting neutrophil effector functions, including production of reactive oxygen species (ROS), release of cytotoxic enzyme via degranulation, and neutrophil extracellular traps (NETs) as promising therapeutic strategies for improving atherosclerotic burden and especially plaque stability at the preclinical level.^4, 5^ While the molecular machinery responsible for these neutrophil functions is becoming more clearly characterized, the upstream signaling pathways and transcriptional regulators for these processes in neutrophils are poorly understood.

Signal transducer and activator of transcription 4 (STAT4) is a member of the STAT family of transcription factors. Upstream signaling via IL-12 activates the IL-12 receptor, causing the autophosphorylation of intermediate Janus Associated Kinase 2 (JAK2), which then phosphorylates STAT4 and initiates its dimerization.^6^ Dimerized phospho-STAT4 undergoes nuclear translocation and initiates transcription of various target genes. STAT4 has been well characterized as an essential mediator of T helper 1 (Th1) cell differentiation during the type 1 interferon response.^6^ We have previously established a role for STAT4 in atherosclerosis in mice, whereby global STAT4 deficiency improves atherosclerotic burden primarily associated with altered T follicular helper (Tfh) differentiation and increased presence of protective CD8+ T regulatory cells.^7^ More recently, we have demonstrated that STAT4 is expressed and readily activated in neutrophils and is required for host defense against methicillin-resistant *Staphylococcus aureus* via several STAT4-dependent neutrophil defense mechanisms including chemotaxis, ROS production, and NET formation.^8^ We therefore hypothesized that STAT4 in neutrophils would participate in atherosclerosis development and plaque instability. Here, we demonstrate that neutrophil-specific STAT4 deficiency improves atherosclerotic burden and plaque stability via reductions in neutrophilia, basal neutrophil activation, and *in vivo* migration to atherosclerotic plaques.

## Materials and Methods

### Animals and study design

Animal protocols were approved by the Eastern Virginia Medical School Institutional Animal Care and Use Committee and were cared for according to the National Institutes of Health Guidelines for Laboratory Animal Care and Use. Animals were kept on a 12-hour light/dark cycle with *ad libitum* access to food and water under pathogen-free conditions. *Stat4*^*fl/fl*^ were generated previously^8^ and crossed with *Ldlr*^*-/-*^ mice (JAX, Cat:#002207) and either with *B6*.*129P2-Lyz2tm1(cre)Ifo/J* or B6.Cg-Tg(S100A8-cre,-EGFP)1Ilw/J transgenic mice (JAX, Cat:#004781; Cat:#021614, respectively) to generate control *Stat4*^*fl/fl*^*Ldlr*^*-/*-^, myeloid-specific *Stat4*^*ΔLysM*^*Ldlr*^*-/-*^ and neutrophil-specific *Stat4*^*ΔS100A8*^*Ldlr*^*-/-*^ *Stat4*-deficient *Ldlr*^*-/-*^ mice. To induce chronic hypercholesterolemia and advanced atherosclerosis, 8-12-week-old male mice were placed on a high fat diet with 0.15% added cholesterol (HFD-C, fat: 60% kcal; carbohydrate: 26% kcal, Bio-Serv F3282) for 28 weeks until sacrifice.

### Quantification of atherosclerosis

Formalin-fixed, paraffin embedded hearts were serial-sectioned at 5 µm thickness from the base of the aortic root at the point of first appearance of the aortic valve leaflets until their disappearance. Three sections equidistant (100 µm) from the midpoint of the valve were used for analysis. Modified Russel-Movat pentachrome staining was performed using a commercially available kit (ScyTek Laboratories). Total plaque area, necrotic core area (% of total plaque area), and fibrotic cap area (% of total plaque area) were measured using ImageJ software (National Institutes of Health).

### Immunofluorescence of fibrotic cap smooth muscle cells

Aortic root paraffin sections were deparaffinized and heat-mediated antigen retrieval was performed using Tris/EDTA (10 mM Tris, 1 mM EDTA, 0.05% Tween, pH 9.0). Sections were blocked for 1 hour at RT with 2% BSA in PBS, followed by incubation with anti-αSMA-Alexa Fluor 647 primary antibody at 4°C. After extensive washing in PBS + 0.1% Tween, sections were counterstained for 5 min with 1µg/mL DAPI (Sigma) and mounted with AquaMount (Polysciences, Inc.). Samples were visualized using a Zeiss AX10 confocal microscope. Fibrous cap smooth muscle cells were counted by positive αSMA staining and normalized to fibrous cap area measured using Zen Black image processing software (Zeiss).

### NanoString gene expression analysis

Circulating neutrophils from chow-fed *Stat4*^*fl/fl*^*Ldlr*^*-/-*^ and *Stat4*^*ΔLysM*^*Ldlr*^*-/-*^, and HFD-C-fed *Stat4*^fl/fl^*Ldlr*^*-/-*^ and *Stat4*^*ΔLysM*^*Ldlr*^*-/-*^ mice were isolated using negative selection (StemCell) and immediately resuspended in lysis buffer (Buffer RLT, Qiagen) supplemented with 1% 2-mercaptoethanol for RNase inhibition. Cells from two mice were pooled for each sample, with n=3 for each group. Neutrophil purity was determined to be >90% pure via flow cytometry. RNA isolation was performed using RNeasy Mini Kit (Qiagen) and was quantified using a Qubit 4 fluorometer (ThermoFisher). Total RNA (25 ng) was assayed using the nCounter® PanCancer Immune Profiling Panel according to the manufacturer’s protocol.

Data was imported and analyzed using ROSALIND®, with a HyperScale architecture developed by ROSALIND, Inc. (San Diego, CA). Normalization, fold changes and p-values were calculated using criteria provided by Nanostring. ROSALIND® follows the nCounter® Advanced Analysis protocol of dividing counts within a lane by the geometric mean of the normalizer probes from the same lane. Housekeeping probes to be used for normalization are selected based on the geNorm algorithm as implemented in the NormqPCR R library. Differentially expressed genes were determined by >1.5-fold change in Log2 normalized expression with a false discovery rate below 0.05.

### Blood hematology

Following euthanasia by CO2 asphyxiation, blood was collected via cardiac puncture and transferred to 1.5 mL microcentrifuge tubes containing 10 µL K2EDTA as an anticoagulant. Blood was collected from the mice at the same time each day to account for circadian variation in neutrophil abundance.^9^ Hematology analysis was performed using a VetScan HM5C hematology analyzer (Abaxis). Plasma total cholesterol was measured using the Cholesterol E method (FUJIFILM Wako Pure Chemical Corporation).

### Flow cytometry

Erythrocytes from 100 µL whole blood were lysed via 10 min ACK (150mM NH4Cl, 10mM KHCO3, 0.1 mM Na2EDTA, pH 7.2-7.4) lysis on ice, followed by Fc blocking with 2% normal rat serum for 10 mins. Cells were stained for 30 mins at 4°C with flow cytometry antibodies as indicated in Supplemental Table 1. For hematopoiesis experiments, bone marrow was collected by flushing tissue from hind tibias and femurs with cold PBS using a 25-gauge needle and syringe, followed by 5 mins ACK lysis. Data were acquired using a 4-laser Aurora spectral flow cytometer (Cytek Inc). Raw data were unmixed using SpectroFlo (Cytek) and analyzed with FlowJo V10 software (BD Biosciences).

### In vivo migration

Bone marrow from 28-week HFD-C fed *Stat4*^*fl/fl*^*Ldlr*^*-/-*^ and *Stat4*^*ΔLysM*^*Ldlr*^*-/-*^ was isolated, lysed of erythrocytes, and labeled with CellTrace Far Red and CellTrace Violet, respectively. 10 × 10^6^ cells of each genotype were combined in a 1:1 ratio and adoptively transferred into aged, atherosclerotic *Apoe*^*-/-*^ mice. 16 hrs later, aortas from the recipient mice were dissected, copiously perfused with PBS, enzymatically digested,^10^ and analyzed for content of labeled neutrophils by flow cytometry. Neutrophil frequency was normalized to the exact proportion of donor *Stat4*^*fl/fl*^*Ldlr*^*-/-*^ to *Stat4*^*ΔLysM*^*Ldlr*^*-/-*^ cells from the starting population, and presented as fold difference in frequency of donor-derived, CD45+CD11b+Ly6G+ *Stat4*^*ΔLysM*^*Ldlr*^*-/-*^ neutrophils compared to *Stat4*^*fl/fl*^*Ldlr*^*-/-*^ neutrophils from each digested recipient aorta.

### Statistical analysis

Data are represented as mean±SEM. Outliers were determined by mean±1.5 x SD and were omitted if these criteria were met. Statistical analysis was performed using Prism 9.4.1 (GraphPad). Normality was determined by Shapiro-Wilk test, and comparisons between groups were performed by unpaired Student t test for experiments with 2 groups, and by one-way ANOVA followed by Tukey multiple comparisons for data with >2 groups.

## Results

### Myeloid and neutrophil-specific Stat4 deficiency improves advanced atherosclerosis and promotes plaque stability

As advanced atherosclerotic plaques develop, not only do they increase in size but also develop features of instability which can lead to rupture and downstream MI or stroke. Necrotic cores are acellular zones that form in part from defective clearance of apoptotic and/or necroptotic cells by macrophages and are typically located near the medial/intimal border. Additionally, thinning of the protective fibrous cap on the plaque luminal surface represents another common feature of plaque destabilization. As macrophages and neutrophils play key roles in both processes,^11^ we asked whether STAT4 deficiency in these cells during advanced atherosclerosis would improve either of these destabilizing features within the aortic root. To examine the role of STAT4 in myeloid cells during advanced atherosclerosis, we successfully generated myeloid-specific (*Stat4*^*ΔLysM*^*Ldlr*^*-/-*^), neutrophil-specific (*Stat4*^*ΔS100A8*^ *Ldlr*^*-/-*^), and *Stat4*-sufficient floxed control (*Stat4*^fl/fl^*Ldlr*^*-/-*^) mice. Following 28 weeks of HFD-C feeding, myeloid and neutrophil STAT4-deficiency resulted in reduced atherosclerotic plaque burden, as well as improvements in plaque stability by decreased necrotic core ratio and increased fibrous cap ratio (Fig.1A-D). This reduction was independent from total plasma cholesterol or degree of weight gain (Supplemental Figure 1). As VSMCs play a pivotal role in maintaining the integrity of the fibrotic cap,^12^ we investigated the abundance of VSMCs within the fibrotic cap by immunohistochemistry and detected an increase in fibrous cap VSMCs in both *Stat4*^*ΔLysM*^*Ldlr*^*-/-*^ and *Stat4*^*ΔS100A8*^ *Ldlr*^*-/-*^ mice compared to *Stat4*^*fl/fl*^*Ldlr*^*-/-*^ controls (Fig.1E). Despite a putative proinflammatory role of STAT4 in monocytes and macrophages,^13^ *Stat4*^*ΔLysM*^*Ldlr*^*-/-*^ showed no additional improvements in plaque burden nor plaque stability compared to *Stat4*^*ΔS100A8*^ *Ldlr*^*-/-*^ mice. These data indicate that STAT4 may play a more dominant proinflammatory role in neutrophils compared to other myeloid cells during advanced atherosclerosis. Due to unforeseen breeding difficulties in *Stat4*^*ΔS100A8*^ *Ldlr*^*-/-*^ mice, the remaining in vitro analyses were conducted from *Stat4*^*ΔLysM*^*Ldlr*^*-/-*^ mice.

**Figure 1.**
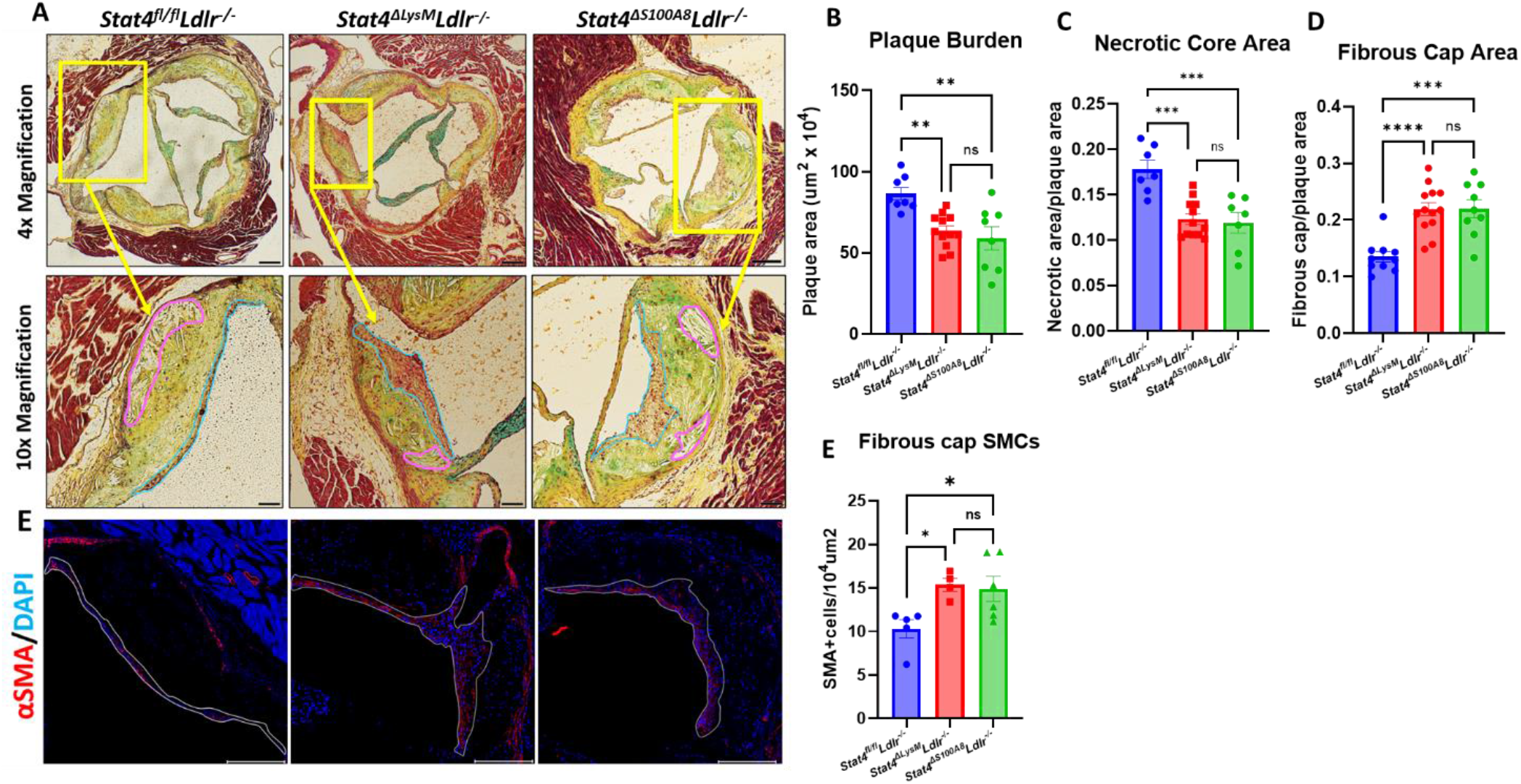
Myeloid and neutrophil-specific STAT4 deficiency improves advanced atherosclerosis and promotes plaque stability. (A) Representative 4X (scale bar = 100 µm) and 10X (scale bar = 200 µm) images of Modified Russel-Movat stained aortic sinus sections, with necrotic core and fibrous cap regions drawn in pink and blue, respectively. (B-D) ImageJ quantifications of total plaque burden, necrotic core area, and fibrous cap area. (E) Representative 10x images of plaques stained for α-SMA with DAPI counterstain (scale bar=200 µm) and (F) quantification of α-SMA+ cells per 10^4^ um^2^. Quantifications represent the average of three equidistant sections per animal. Data represented as mean±SEM. One-way ANOVA with Tukey’s multiple comparisons. n=8-11 mice/group for Movat analysis. n=4-6 mice/group for immunofluorescence. *p<0.05, **p<0.01, ***p<0.001. Two independent experiments.

### Stat4 deficiency reduces circulating neutrophils via impaired medullary myelopoiesis

To assess the role of STAT4 in neutrophil biology during advanced atherosclerosis, we first sought to determine whether STAT4-deficiency affects circulating neutrophil abundance. Indeed, we detected reduced frequency and total circulating numbers of neutrophils (Fig.1A-B), as well as reduced neutrophil-to-lymphocyte ratio in *Stat4*^*ΔLysM*^*Ldlr*^*-/-*^ mice compared to *Stat4*^*fl/fl*^*Ldlr*^*-/-*^ control mice (Fig.2C). Lymphocyte frequency (%) was increased in the *Stat4*^*ΔLysM*^*Ldlr*^*-/-*^ mice, which is likely due to the drop in neutrophil frequency (Fig.2G). Meanwhile, total leukocyte, lymphocyte, and monocyte absolute counts remained unaffected (Fig.2).

**Figure 2.**
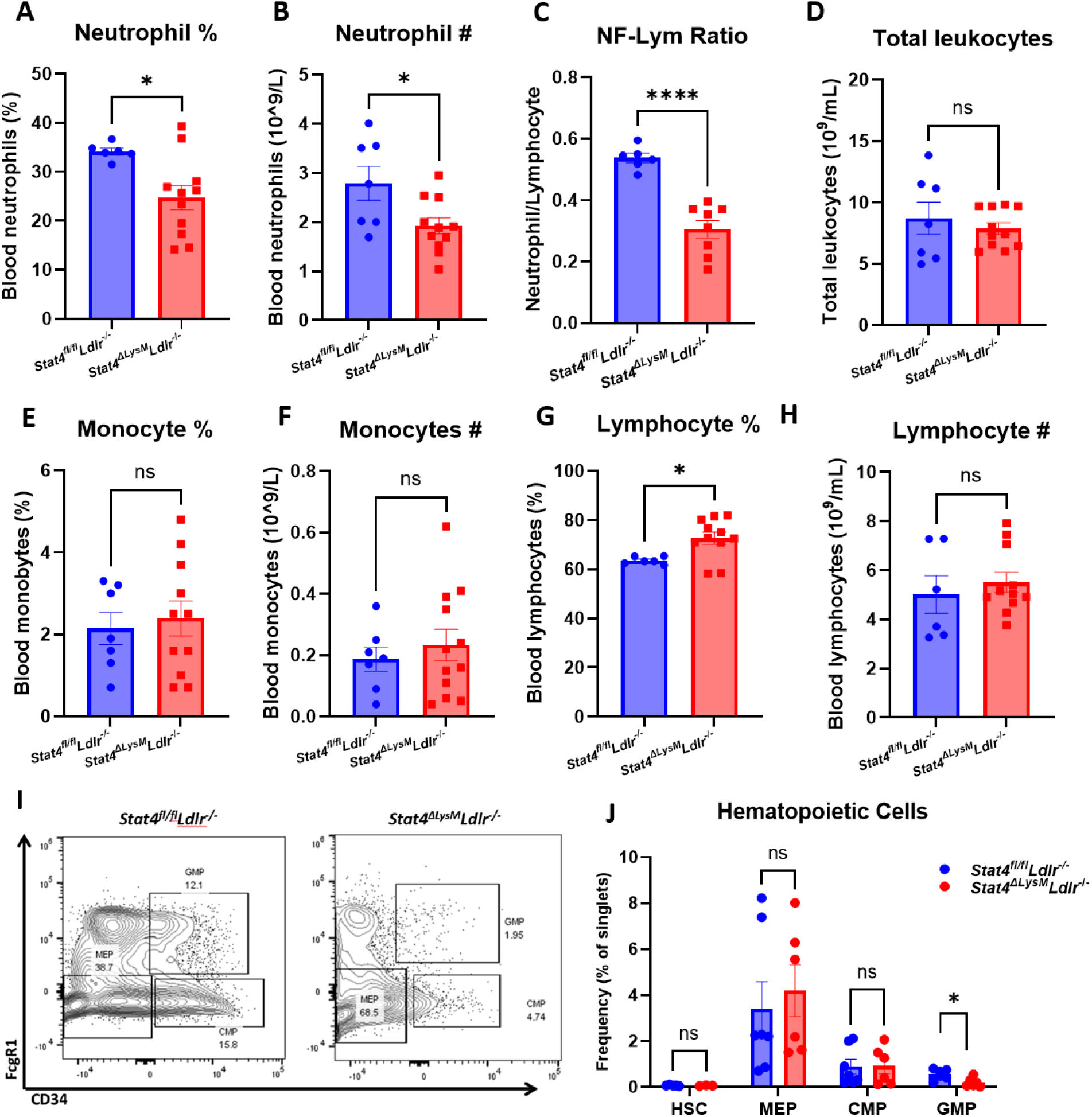
Myeloid *Stat4* deficiency reduces circulating neutrophils via impaired medullary myelopoiesis. (A-H) VetScan automated hematology data showing reduced abundance of circulating neutrophils in *Stat4*^*ΔLysM*^*Ldlr*^*-/-*^ *vs Stat4*^fl/fl^*Ldlr*^*-/-*^ mice. (I) Representative FACS plots for identifying granulocyte-macrophate progenitor (GMP), common myeloid progenitor (CMP), and megakaryocyte-erythrocyte progenitor (MEP) populations from Lin^-^Sca1^-^cKit^+^ bone marrow cells. (J) Quantification of hematopoietic populations as a percentage of total singlets. Data represented as mean±SEM. Unpaired student’s t test. n=6-12 mice/group. *p<0.05, ****p<0.0001. 2 independent experiments.

Neutrophilia associated with advanced atherosclerosis arises from chronic dyslipidemia and influences hyperactivated myelopoiesis^14^. The classical model of medullary myelopoiesis begins when hematopoietic stem cells (HSCs) differentiate into common myeloid progenitors (CMPs), followed by transition to granulocyte-macrophage progenitors (GMPs), which can then give rise to immature and eventually mature neutrophils.^15^ To determine whether the reduction in neutrophilia in STAT4-deficient mice resulted from impaired myelopoiesis, bone marrow from *Stat4*^*ΔLysM*^*Ldlr*^*-/-*^ and *Stat4*^*fl/fl*^*Ldlr*^*-/-*^ mice were analyzed for major hematopoietic cell populations. Interestingly, we found reduced frequency of GMPs (Lin^-^Sca1^-^cKit^+^CD34^+^FcγR1^+^) in the *Stat4*^*ΔLysM*^*Ldlr*^*-/-*^ compared to *Stat4*^*fl/fl*^*Ldlr*^*-/-*^ mice (Fig.2I-J). However, frequencies of HSCs, megakaryocyte-erythrocyte progenitors ((Lin^-^Sca1^-^cKit^+^CD34^-^FcγR1^-^), and CMPs ((Lin^-^ Sca1^-^cKit^+^CD34^+^FcγR1^-^) remained unchanged. Together, these findings suggest that myeloid STAT4 plays a role in neutrophil differentiation during hyperlipidemia-induced myelopoiesis and contributes to the circulating pool of neutrophils during advanced atherosclerosis.

### Neutrophil transcriptome is modified during advanced atherosclerosis and partially modulated by Stat4

Since STAT4 functions as a transcription factor, it was important to address to what degree *Stat4* influences the neutrophil gene expression profile during atherosclerosis. To accomplish this, we isolated blood neutrophils from 28-week HFD-C-fed *Stat4*^fl/fl^*Ldlr*^*-/-*^ and *Stat4*^*ΔLysM*^*Ldlr*^*-/-*^ mice, as well as age matched, chow-fed *Stat4*^fl/fl^*Ldlr*^*-/-*^ and *Stat4*^*ΔLysM*^*Ldlr*^*-/-*^ neutrophils and analyzed an array of 750 genes related to innate immunity using the NanoString nCounter PanCancer Immune Profiling panel. We found 77 differentially upregulated genes from HFD-C fed *Stat4*^flox/flox^*Ldlr*^*-/-*^ neutrophils compared to chow-fed controls (Fig.3A). Many of the most upregulated genes have known proatherogenic implications, including, *S100A8*^*16*^, *Cxcl2*^*17*^, *Lcn2*^*18, 19*^, *Camp*^*20*^, and several others. S100A8, a secretory alarmin produced by all myeloid cells, acts as a secretory signaling molecule released in response to inflammatory stimuli in neutrophils. Several studies indicate a direct role for S100A8 in vascular inflammation via chemoattractant properties inducing migration of other phagocytes and VCAM-1/ICAM-1 inducing activation of endothelial cells (ECs), and stimulating the differentiation of myeloid progenitor cells with the release of Ly6C^hi^ bone marrow monocytes.^16^

**Figure 3.**
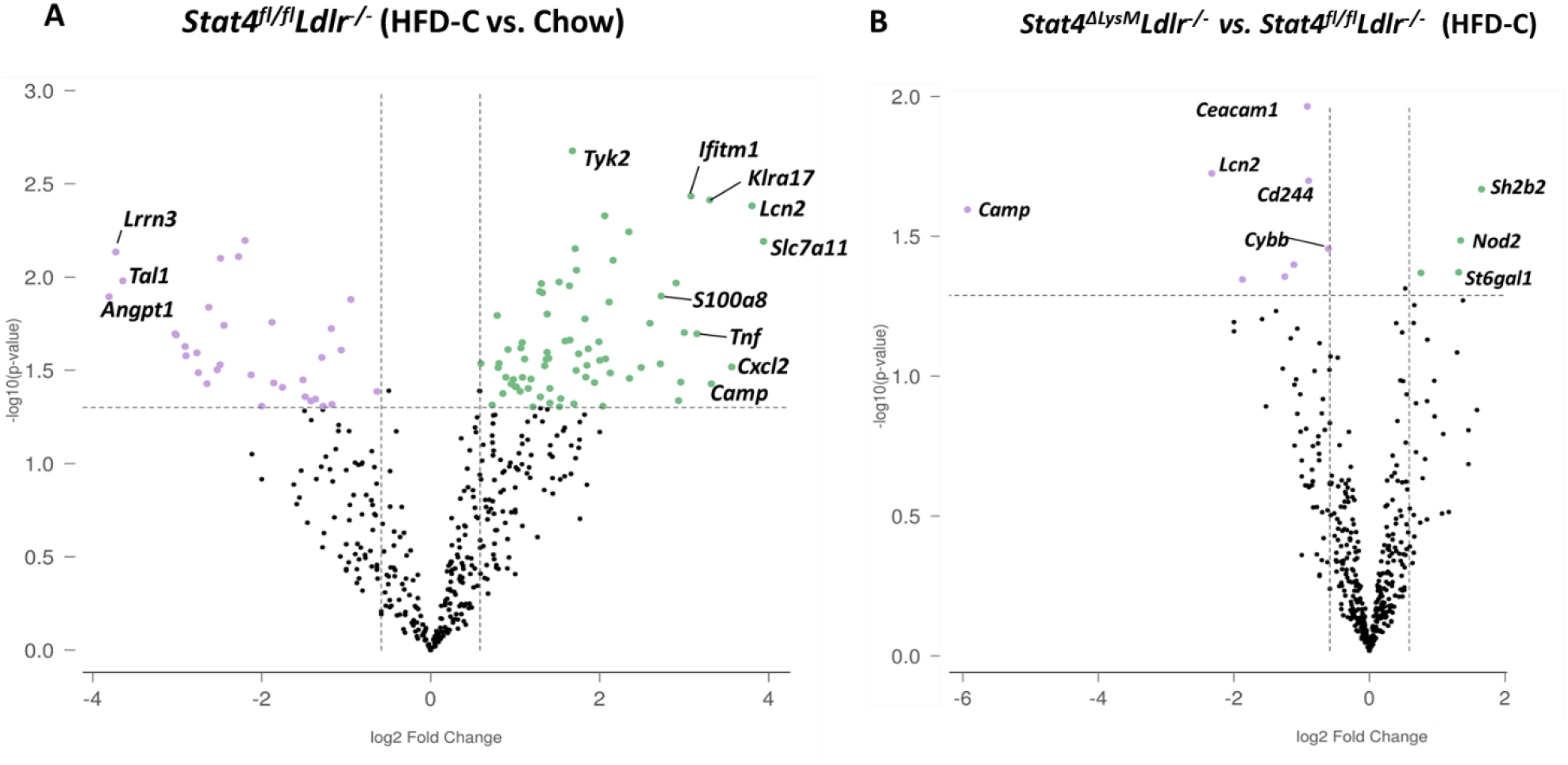
Neutrophil transcriptome is enhanced during advanced atherosclerosis and partially modulated by *Stat4* deficiency. (A) Volcano plot from Nanostring gene expression data showing upregulated genes from isolated blood neutrophils of 28-week HFD-C fed *Stat4*^*fl/fl*^*Ldlr*^*-/-*^ mice (green) compared to age-matched chow-fed controls (purple). (B) Volcano plot showing differentially downregulated genes in *Stat4*^*ΔLysM*^*Ldlr*^*-/-*^ neutrophils (purple) compared to *Stat4*^*fl/fl*^*Ldlr*^*-/-*^ neutrophils (green). Differentially expressed genes were determined as >1.5-fold difference with adjusted p value <0.05. n=3 mice per group, with neutrophils pooled from 2 mice per sample.

Next, we sought to determine how STAT4 deficiency in neutrophils affects their expression profile upon atherosclerosis development. When comparing HFD-C fed *Stat4*^*ΔLysM*^*Ldlr*^*-/-*^ to *Stat4*^*fl/fl*^*Ldlr*^*-/-*^ neutrophils, we found only 9 differentially downregulated genes (Fig.3B). However, several of these genes were among the most highly upregulated in *Stat4*^*fl/fl*^*Ldlr*^*-/-*^ neutrophils compared to chow controls, including *Camp* and *Lcn2*. In addition, we found a downregulation of *Cybb*, which encodes for NADPH oxidase-2 (NOX-2). NOX-2 is critical for production of superoxide species required for antimicrobial killing mechanism in neutrophils, but its upregulation in atherosclerotic plaques is well known to contribute directly to plaque development and instability.^21, 22^ Together, these data emphasize the milieu of inflammatory genes upregulated in neutrophils during atherosclerosis and highlight STAT4 as a novel regulator of several key proatherogenic genes in neutrophils.

### Stat4 deficiency impairs mitochondrial superoxide production, degranulation, and in vivo migration in neutrophils

To further our understanding of how STAT4 contributes to neutrophil biology during advanced atherosclerosis at the functional level, we analyzed several aspects of neutrophil activation. To corroborate our finding of reduced *Cybb* expression, we sought to determine whether *Stat4*^*ΔLysM*^*Ldlr*^*-/-*^ circulating neutrophils produce less basal superoxide compared to *Stat4*^fl/fl^*Ldlr*^*-/-*^ neutrophils. Indeed, we detected a reduction of CD11b+Ly6G+ neutrophils positive for MitoSOX, which stains specifically for mitochondrial superoxides (Fig.4A-B). Another mechanism of antimicrobial neutrophil defense is the release of intracellular granules containing antibacterial enzymes; this process becomes dysregulated during atherosclerosis and contributes to disease progression and plaque vulnerability.^23, 24^ CD63 is expressed on the surface of intracellular neutrophil granules and is present on the plasma membrane during degranulation.^25^ Interestingly, we detected a reduction in surface CD63+ neutrophils from *Stat4*^*ΔLysM*^*Ldlr*^*-/-*^ mice, suggesting that STAT4 may play a role in neutrophil activation and release of cytotoxic granules (Fig.4C-D). Finally, neutrophil-platelet interactions and aggregates are elevated during atherosclerosis and contribute to atheroprogression via the promotion of neutrophil recruitment to the plaque and increasing the generation of thrombi.^26^ Detection of these aggregates can be measured by the presence of platelet marker CD41 on the surface of neutrophils. We uncovered a significant reduction in the frequency of CD41+CD11b+Ly6G+ neutrophils from *Stat4*^*ΔLysM*^*Ldlr*^*-/-*^ mice compared to *Stat4*^*fl/fl*^*Ldlr*^*-/-*^ controls (Fig.4E-F) Together, these data demonstrate that STAT4 deficiency results in the reduction of several key aspects of neutrophil activation that impact the progression of atherosclerosis and promote plaque instability.

**Figure 4.**
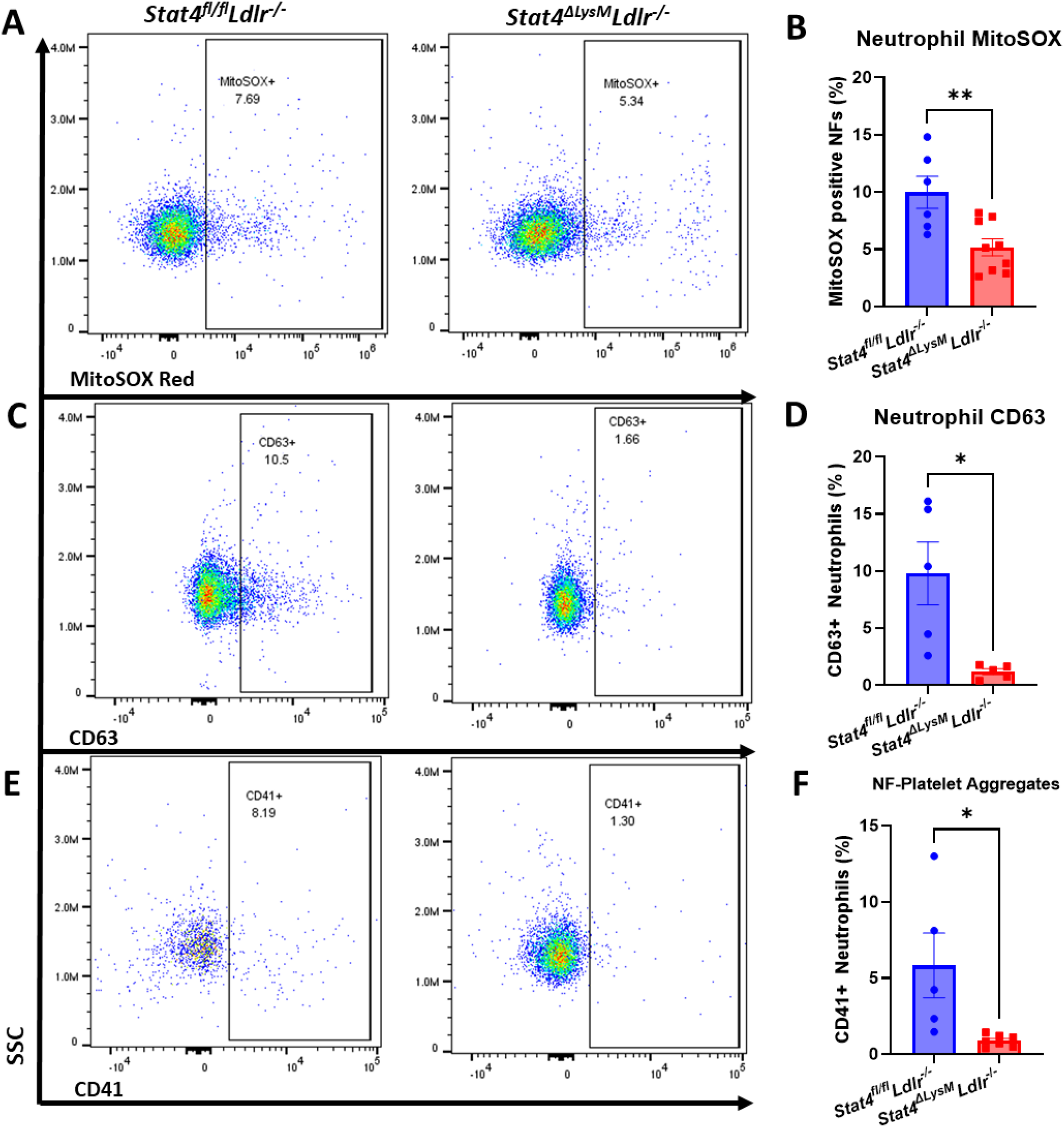
Myeloid STAT4 deficiency reduces several aspects of basal neutrophil activation. (A-B) Representative FACS plots and quantification of basal mitochondrial superoxide production by MitoSOX Red positive stained blood neutrophils. (C-D) Representative FACS plot and quantification of surface expression of neutrophil degranulation marker CD63. (E-F) Representative FACS plots and quantification of neutrophil-platelet aggregates by presence of platelet marker CD41 staining on neutrophils. All plots represent CD11b+Ly6G+ blood neutrophils. Data represent mean±SEM. Unpaired student’s t test. n= 5-9 mice/group. *p<0.05, **p<0.01. 2-3 independent experiments.

Neutrophils are among the most short-lived immune cells and are continuously replenished into the circulation from the bone marrow. The migration of neutrophils to atherosclerotic plaques is a key area of study devoted to mechanisms of development of plaque instability. Indeed, a crucial role of activated neutrophils in the formation of unstable plaques via uncontrolled release of ROS, cytotoxic granules, matrix degrading enzymes, etc., has been demonstrated in several papers.^27^ However, very little is known about mechanisms of neutrophil recruitment into the atherosclerotic aorta. To address the role of STAT4 in neutrophil migration to sites of atherosclerosis, we analyzed several aspects of neutrophil migration both *in vitro* and *in vivo*. As CCR1 and CCR2 serve as key chemokine receptors for neutrophils and respecting chemokines are highly expressed within the aortic wall,^28, 29^ we analyzed expression of these markers on the surface of circulating neutrophils. Firstly, we discovered reduced expression of chemokine receptors 1 and 2 (CCR1, CCR2) on the surface of circulating *Stat4*^*ΔLysM*^*Ldlr*^*-/-*^ neutrophils compared to *Stat4*^*fl/fl*^*Ldlr*^*-/-*^ controls (Fig.5A-B). To address whether these differences in CCR1 and CCR2 expression impair neutrophil migration *in vivo*, we prelabeled *Stat4*^*ΔLysM*^*Ldlr*^*-/-*^ and *Stat4*^*fl/fl*^*Ldlr*^*-/-*^ bone marrow neutrophils with cell tracing dyes CellTrace Violet and CellTrace Far Red, respectively, followed by intravenous adoptive transfer of a 1:1 ratio of these cells into aged, atherosclerotic *Apoe*^-/-^ mice.^30^

**Figure 5.**
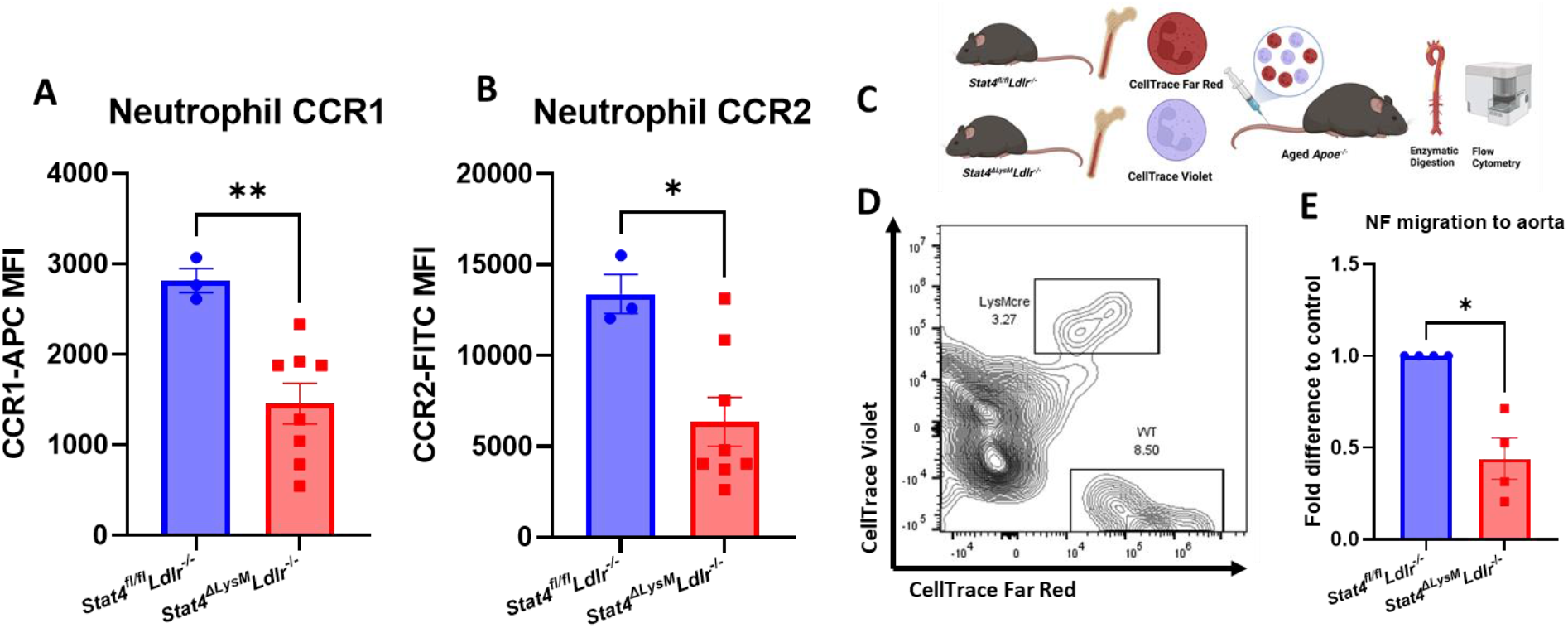
*Stat4-*deficient neutrophils have reduced chemokine receptor expression and impaired *in vivo* migration to atherosclerotic aorta. (A-B) Quantification of mean fluorescence intensity for chemokine receptors 1 and 2 expression on blood neutrophils. (C) Schematic of workflow for competitive homing of prelabeled, adoptively transferred *Stat4*^*fl/fl*^*Ldlr*^*-/-*^ and *Stat4*^*ΔLysM*^*Ldlr*^*-/-*^ bone marrow cells into aged, atherosclerotic *Apoe*^*-/-*^ mice followed by enzymatic digestion of recipient aortas for FACS analysis of donor neutrophil homing to aorta. (D) Representative FACS plot and (E) Fold difference of adoptively transferred CD11b+Ly6G+ neutrophils within the aorta of recipient mice, normalized to distribution of starting population. (A-B) Data represent mean±SEM. Unpaired Student’s t test. n=3-8. * p<0.05, **p<0.01. 2 independent experiments.

Following 16 hours post-transfer, aortas from recipient mice were dissected, enzymatically digested, and analyzed for abundance of donor CD45+CD11b+Ly6G+ neutrophils (Fig.5C). We established a recruitment rate of STAT4-sufficient *Stat4*^*fl/fl*^*Ldlr*^*-/-*^ neutrophils as 1 and calculated a relative ratio of emigrated *Stat4*^*ΔLysM*^*Ldlr*^*-/-*^ neutrophils to the aorta for each day of independent experiments. Overall, we detected a reduction in donor-derived *Stat4*^*ΔLysM*^*Ldlr*^*-/-*^ neutrophils within the recipient aortas (Fig.5D-E), suggesting that STAT4-deficient neutrophils have reduced capacity to migrate into the atherosclerotic vessels. Altogether, these findings demonstrate STAT4 as a novel regulator of neutrophil homing to sites of atherosclerosis.

## Discussion

In this present study, we have increased our understanding of the proatherogenic role of neutrophils by revealing several STAT4-dependent mechanisms that contribute to atherosclerotic plaque progression and instability. Generation of both myeloid-specific and neutrophil-specific STAT4-deficient mice allowed us to pinpoint to what degree STAT4 contributes to atherosclerosis within the entire myeloid compartment and individually from neutrophils. The *LysM-*cre transgenic mouse has become firmly established as the premier model for achieving myeloid-specific transgenic mice when crossed with a floxed mouse on either the *Ldlr*^*-/-*^ or *Apoe*^*-/-*^ atherogenic backgrounds with Cre-negative floxed mice serving as appropriate controls.^31-36^ In addition, *LysM*cre-*Ldlr*^*-/-*^ mice fed high fat diet for either 3 or 6 months develop considerable atherosclerosis, thus demonstrating that the cre recombinase expression alone does not affect the development of atherosclerosis.^37^ Interestingly, despite the disruption of *Lyz2* expression from the insertion of the cre recombinase at the first coting ATG of the *Lyz2* gene, absence of *Lyz2* shows no effect on either monocyte or neutrophil function in a model of sterile inflammation.^38^ We were indeed surprised to discover that STAT4-deficiency in neutrophils alone achieved the same level of reduced plaque burden and improved plaque stability compared to STAT4 deficiency within all myeloid cells. Additionally, our finding that myeloid STAT4 deficiency reduced the abundance of GMPs in the bone marrow, leading to a decrease in circulating neutrophil abundance while leaving monocytes unaffected prompted us to shift our focus more towards the effect of STAT4 deficiency on neutrophils during atherosclerosis. The STAT4-dependent decrease in circulating neutrophils and associated decrease in neutrophil-to-lymphocyte ratio is of substantial clinical significance. Not only does neutrophil-to-lymphocyte ratio associate with increased risk of death, myocardial infarction, and coronary heart disease,^39^ but has recently been strongly correlated with carotid artery plaque instability.^40^ During granulopoiesis, several maturation steps occur between the GMP and mature neutrophil. This includes the progression from myeloblast to promyelocyte, myelocyte, metamyelocyte, band cell, and finally the mature neutrophil released into circulation.^41^ New developments of neutrophil maturation in the bone marrow have identified several intermediate cell types between the GMP and the mature neutrophil, which include the Ly6B+ neutrophil precursor.^42^ How STAT4 influences any or each of these maturation steps during granulopoiesis warrants further investigation.

Our investigation into the neutrophil gene expression landscape using the Nanosting nCounter platform adds to the existing body of knowledge that neutrophils upregulate a plethora of proinflammatory genes throughout the process of atherosclerosis. The differential upregulation of only 77 genes in isolated neutrophils from nonatherosclerotic chow-fed *Stat4*^*fl/fl*^*Ldlr*^*-/-*^ compared to atherosclerotic HFD-C fed *Stat4*^*fl/fl*^*Ldlr*^*-/-*^ mice was fewer than expected. Neutrophils have lower RNA content compared to other leukocytes and abundance of RNases are relatively high, and even with RNase inhibition during sample preparation, obtaining sufficient quantity and quality of RNA from neutrophils remains a significant challenge.^43^ This could also serve as an explanation to the relatively few downregulated genes in neutrophils from HFD-C fed *Stat4*^*ΔLysM*^*Ldlr*^*-/-*^ compared to *Stat4*^*fl/fl*^*Ldlr*^*-/-*^ mice. Regardless, the assay revealed that STAT4 deficiency resulted in the downregulation of some of the most highly upregulated genes, namely *Camp* and *Lcn2. Camp* encodes for the secondary granule protein CRAMP, and its inhibition in mouse models has been shown to improve atherosclerosis.^44^ Mechanistically, neutrophil-derived CRAMP/DNA complexes have been shown to promote atherosclerosis via direct activation of plasmacytoid dendritic cells.^45^ Likewise, *Lcn* deficiency in late stage atherosclerotic *Ldlr*^*-/-*^ mice improved plaque stability by decreasing necrotic core size.^46^ Perhaps the most impactful discovery from our Nanostring analysis was the reduced expression of *Cybb* (NADPH oxidase 2) as this gene is critical to several neutrophil functions related to oxidative burst-dependent mechanisms which are often dysregulated during atherosclerosis.^47^ Indeed, our demonstrated correlation of reduced *Cybb* expression with reduced mitochondrial superoxide production in neutrophils revealed STAT4 as a novel contributor to neutrophil oxidative stress during atherosclerosis. Furthermore, NOX-2 has been demonstrated to play a key role in neutrophil-platelet interactions during vascular inflammation.^48^ Neutrophil-platelet interactions are elevated in patients with coronary artery disease^49^ and have been associate with increased risk of thrombosis.^50^ The reduced frequency of neutrophil-platelet aggregates in *Stat4*^*ΔLysM*^*Ldlr*^*-/-*^ mice further supports STAT4 as a critical driver of neutrophil activation during advanced atherosclerosis and the promotion of plaque destabilization. Recently, the cathelicidin LL-37, which is the human homologue to murine CRAMP, was shown to be plentiful in thrombi of patients with acute MI and primes platelets for activation and neutrophil-platelet interactions.^51^ Since our Nanostring data revealed the STAT4-dependent downregulation of *Camp* in *Stat4*^*ΔLysM*^*Ldlr*^*-/-*^ neutrophils, this could be a possible mechanistic explanation for the reduction in circulating neutrophil-platelet interactions and resulting improvement in plaque stability.

Rapid recruitment and migration from circulation to sites of inflammation within vascular structures and other tissues is a hallmark of neutrophil biology.^52^ Neutrophils respond to a complex network of various chemokines that become elevated under atherosclerotic conditions. Several chemokine receptors are required for neutrophil recruitment to atherosclerosis in large arteries, including CCR1 and CCR2.^53^ Furthermore, expression of CCR1/2 further increases upon extravasation, and is critical for its phagocytic activity and ROS production.^54^ Ligands for these receptors include macrophage-derived CCL3 and platelet-derived CCL5, which are critical for neutrophil homing to atherosclerotic plaques.^55^ Invasion of neutrophils into atherosclerotic plaques has been shown to directly influence several features of plaque stability by decreasing plaque collagen content, decreasing fibrous cap thickness, and enlarging the intraplaque necrotic core all within 24 hours of neutrophil challenge.^56^ Here, we demonstrate that STAT4 deficiency in neutrophils results in reduced expression of chemokine receptors while exhibiting reduced capacity of activated neutrophils to migrate to sites of atherosclerosis.

Taken together, this study further emphasizes the critical importance of neutrophils during advanced atherosclerosis with a special emphasis on its contribution to late-stage plaque instability. Recent research efforts have begun to dissect the role of individual receptors, enzymes, and metabolic mediators that contribute to the diverse armament of neutrophil effector functions.^57^ However, limited research has uncovered the transcriptional networks that regulate such functions. Our data clearly demonstrate STAT4 as a novel regulator of neutrophil inflammatory mechanisms during advanced atherosclerosis, which could offer new therapeutic avenues for the prevention of atherosclerotic plaque destabilization.

## Supplementary Material

Refer to Web version on PubMed Central for supplementary material.

## Acknowledgments

This work was supported by NIH R01HL142129 (EVG and JLN), R01HL139000 (EVG), and AHA pre-doctoral fellowship 20PRE35180156 (AKM). Flow cytometry analysis/cell sorting was performed at the EVMS Flow Cytometry Core Facility.”

---

**Table.**
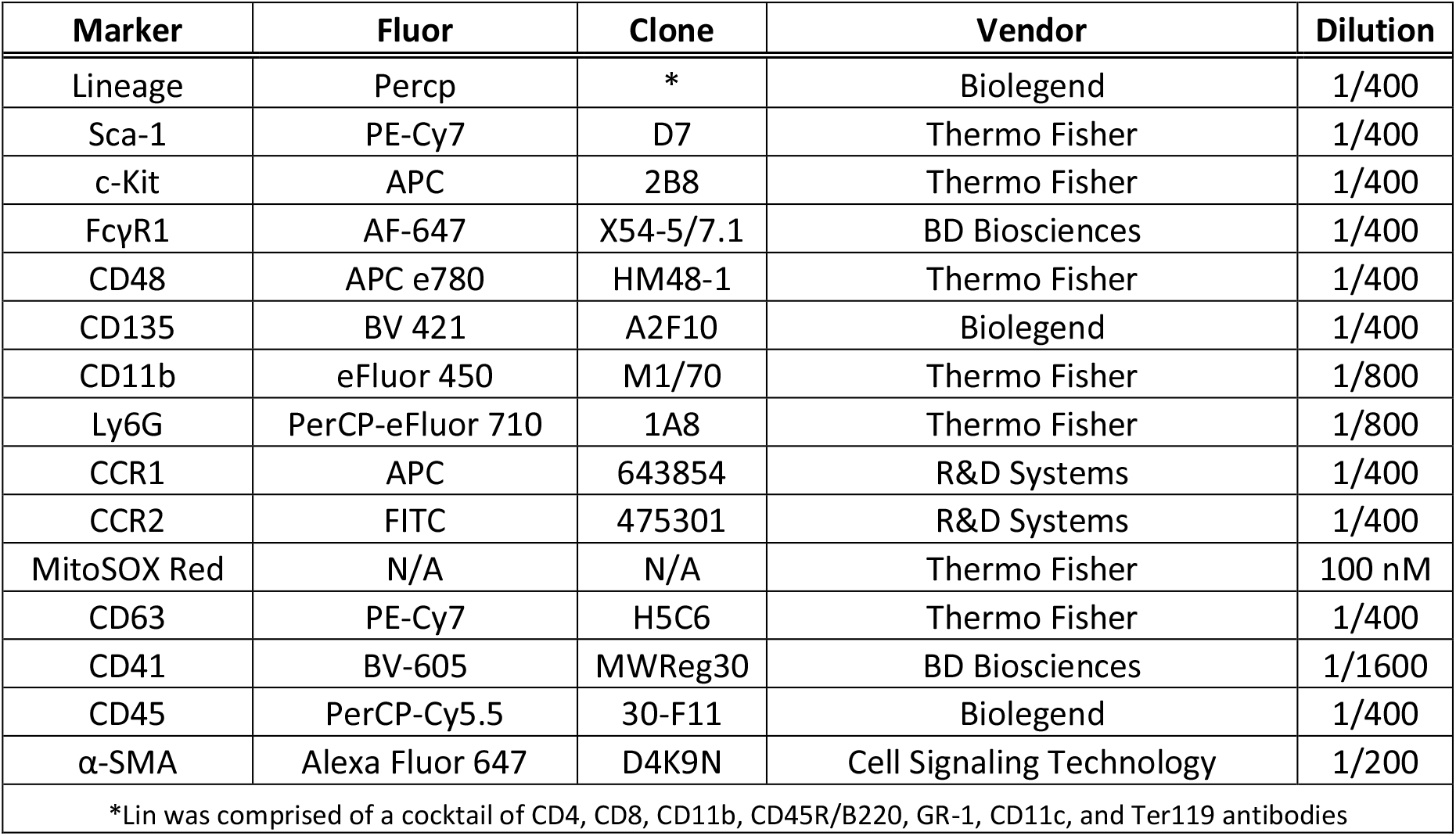
List of all antibodies, dilutions, and vendor sources.

**Supplemental Figure 1.**
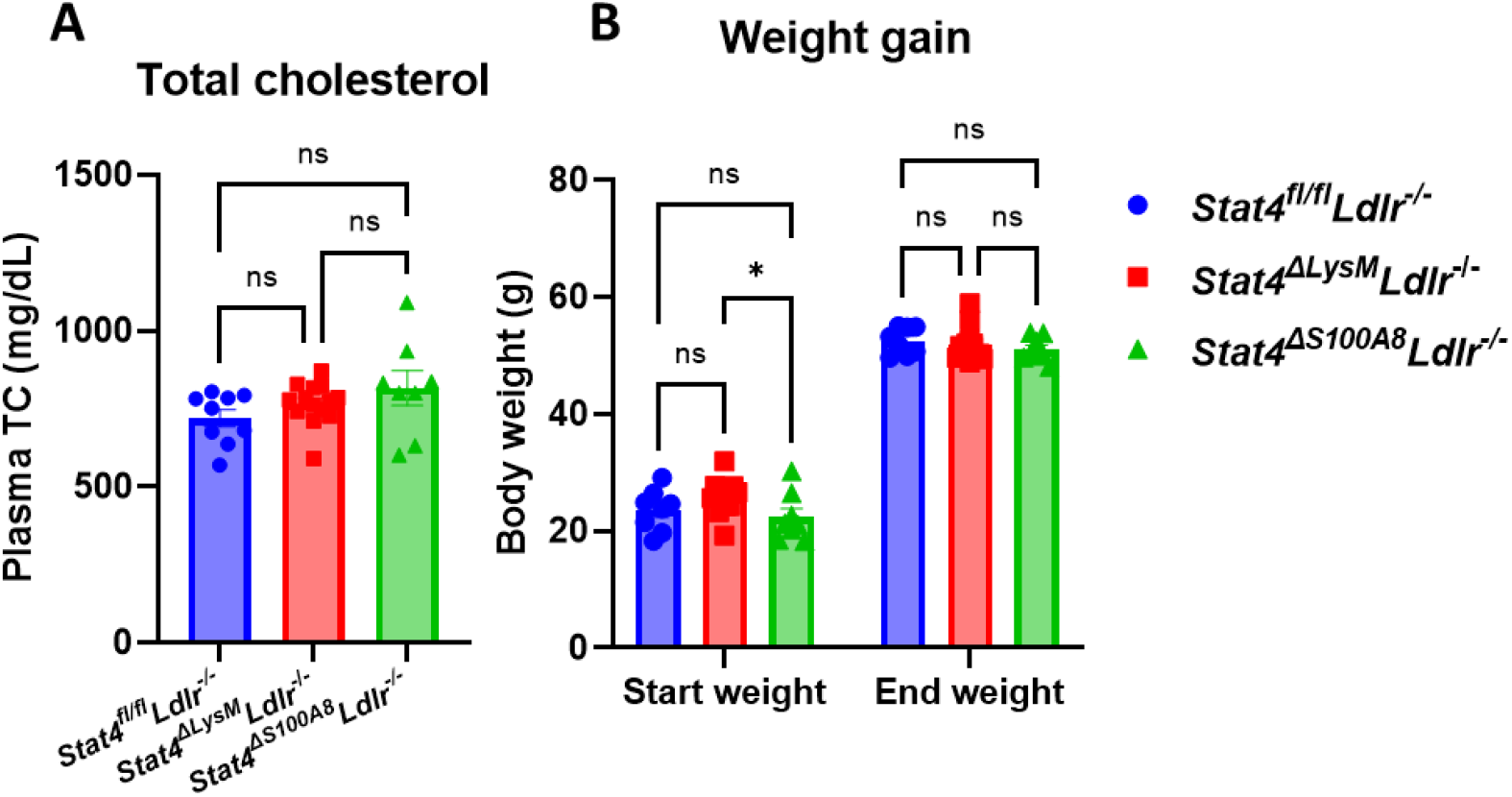
(A) Total plasma cholesterol measurements at 28 weeks HFD-C feeding. (B) Beginning and final body weight measurements at 0 and 28 weeks HFD feeding. Data represent mean±SEM. One-way ANOVA (A) and Two-way ANOVA (B) with Tukey’s multiple comparisons test. *p<0.05.

## References

1. Tsao CW, Aday AW, Almarzooq ZI, Alonso A, Beaton AZ, Bittencourt MS, Boehme AK, Buxton AE, Carson AP, Commodore-Mensah Y, Elkind MSV, Evenson KR, Eze-Nliam C, Ferguson JF, Generoso G, Ho JE, Kalani R, Khan SS, Kissela BM, Knutson KL, Levine DA, Lewis TT, Liu J, Loop MS, Ma J, Mussolino ME, Navaneethan SD, Perak AM, Poudel R, Rezk-Hanna M, Roth GA, Schroeder EB, Shah SH, Thacker EL, VanWagner LB, Virani SS, Voecks JH, Wang NY, Yaffe K and Martin SS. Heart Disease and Stroke Statistics-2022 Update: A Report From the American Heart Association. Circulation. 2022;145:e153–e639.

2. Roy P, Orecchioni M and Ley K. How the immune system shapes atherosclerosis: roles of innate and adaptive immunity. Nat Rev Immunol. 2022;22:251–265.

3. Fernandez DM, Rahman AH, Fernandez NF, Chudnovskiy A, Amir ED, Amadori L, Khan NS, Wong CK, Shamailova R, Hill CA, Wang Z, Remark R, Li JR, Pina C, Faries C, Awad AJ, Moss N, Bjorkegren JLM, Kim-Schulze S, Gnjatic S, Ma’ayan A, Mocco J, Faries P, Merad M and Giannarelli C. Single-cell immune landscape of human atherosclerotic plaques. Nat Med. 2019;25:1576–1588.

4. Silvestre-Roig C, Braster Q, Wichapong K, Lee EY, Teulon JM, Berrebeh N, Winter J, Adrover JM, Santos GS, Froese A, Lemnitzer P, Ortega-Gomez A, Chevre R, Marschner J, Schumski A, Winter C, Perez-Olivares L, Pan C, Paulin N, Schoufour T, Hartwig H, Gonzalez-Ramos S, Kamp F, Megens RTA, Mowen KA, Gunzer M, Maegdefessel L, Hackeng T, Lutgens E, Daemen M, von Blume J, Anders HJ, Nikolaev VO, Pellequer JL, Weber C, Hidalgo A, Nicolaes GAF, Wong GCL and Soehnlein O. Externalized histone H4 orchestrates chronic inflammation by inducing lytic cell death. Nature. 2019;569:236–240.

5. Franck G, Mawson TL, Folco EJ, Molinaro R, Ruvkun V, Engelbertsen D, Liu X, Tesmenitsky Y, Shvartz E, Sukhova GK, Michel JB, Nicoletti A, Lichtman A, Wagner D, Croce KJ and Libby P. Roles of PAD4 and NETosis in Experimental Atherosclerosis and Arterial Injury: Implications for Superficial Erosion. Circ Res. 2018;123:33–42.

6. Nishikomori R, Usui T, Wu CY, Morinobu A, O’Shea JJ and Strober W. Activated STAT4 has an essential role in Th1 differentiation and proliferation that is independent of its role in the maintenance of IL-12R beta 2 chain expression and signaling. J Immunol. 2002;169:4388–98.

7. Taghavie-Moghadam PL, Waseem TC, Hattler J, Glenn LM, Dobrian AD, Kaplan MH, Yang Y, Nurieva R, Nadler JL and Galkina EV. STAT4 Regulates the CD8(+) Regulatory T Cell/T Follicular Helper Cell Axis and Promotes Atherogenesis in Insulin-Resistant Ldlr(-/-) Mice. J Immunol. 2017;199:3453–3465.

8. Mehrpouya-Bahrami P, Moriarty AK, De Melo P, Keeter WC, Alakhras NS, Nelson AS, Hoover M, Barrios MS, Nadler JL, Serezani CH, Kaplan MH and Galkina EV. STAT4 is expressed in neutrophils and promotes antimicrobial immunity. JCI Insight. 2021;6.

9. Adrover JM, Del Fresno C, Crainiciuc G, Cuartero MI, Casanova-Acebes M, Weiss LA, Huerga-Encabo H, Silvestre-Roig C, Rossaint J, Cossio I, Lechuga-Vieco AV, Garcia-Prieto J, Gomez-Parrizas M, Quintana JA, Ballesteros I, Martin-Salamanca S, Aroca-Crevillen A, Chong SZ, Evrard M, Balabanian K, Lopez J, Bidzhekov K, Bachelerie F, Abad-Santos F, Munoz-Calleja C, Zarbock A, Soehnlein O, Weber C, Ng LG, Lopez-Rodriguez C, Sancho D, Moro MA, Ibanez B and Hidalgo A. A Neutrophil Timer Coordinates Immune Defense and Vascular Protection. Immunity. 2019;50:390–402 e10.

10. Gjurich BN, Taghavie-Moghadam PL and Galkina EV. Flow Cytometric Analysis of Immune Cells Within Murine Aorta. Methods Mol Biol. 2015;1339:161–75.

11. Hansson GK, Libby P and Tabas I. Inflammation and plaque vulnerability. J Intern Med. 2015;278:483–93.

12. Harman JL and Jorgensen HF. The role of smooth muscle cells in plaque stability: Therapeutic targeting potential. Br J Pharmacol. 2019;176:3741–3753.

13. Taghavie-Moghadam PL, Gjurich BN, Jabeen R, Krishnamurthy P, Kaplan MH, Dobrian AD, Nadler JL and Galkina EV. STAT4 deficiency reduces the development of atherosclerosis in mice. Atherosclerosis. 2015;243:169–78.

14. Soehnlein O and Swirski FK. Hypercholesterolemia links hematopoiesis with atherosclerosis. Trends Endocrinol Metab. 2013;24:129–36.

15. Abdali A and Marinkovic G. Assessment of medullary and extramedullary myelopoiesis in cardiovascular diseases. Pharmacol Res. 2021;169:105663.

16. Sreejit G, Abdel Latif A, Murphy AJ and Nagareddy PR. Emerging roles of neutrophil-borne S100A8/A9 in cardiovascular inflammation. Pharmacol Res. 2020;161:105212.

17. Yang J, Liu H, Cao Q and Zhong W. Characteristics of CXCL2 expression in coronary atherosclerosis and negative regulation by microRNA-421. J Int Med Res. 2020;48:300060519896150.

18. Gan J, Zheng Y, Yu Q, Zhang Y, Xie W, Shi Y, Yu N, Yan Y, Lin Z and Yang H. Serum Lipocalin-2 Levels Are Increased and Independently Associated With Early-Stage Renal Damage and Carotid Atherosclerotic Plaque in Patients With T2DM. Front Endocrinol (Lausanne). 2022;13:855616.

19. Oberoi R, Bogalle EP, Matthes LA, Schuett H, Koch AK, Grote K, Schieffer B, Schuett J and Luchtefeld M. Lipocalin (LCN) 2 Mediates Pro-Atherosclerotic Processes and Is Elevated in Patients with Coronary Artery Disease. PLoS One. 2015;10:e0137924.

20. Mihailovic PM, Lio WM, Yano J, Zhao X, Zhou J, Chyu KY, Shah PK, Cercek B and Dimayuga PC. The cathelicidin protein CRAMP is a potential atherosclerosis self-antigen in ApoE(-/-) mice. PLoS One. 2017;12:e0187432.

21. Quesada IM, Lucero A, Amaya C, Meijles DN, Cifuentes ME, Pagano PJ and Castro C. Selective inactivation of NADPH oxidase 2 causes regression of vascularization and the size and stability of atherosclerotic plaques. Atherosclerosis. 2015;242:469–75.

22. Violi F, Carnevale R, Loffredo L, Pignatelli P and Gallin JI. NADPH Oxidase-2 and Atherothrombosis: Insight From Chronic Granulomatous Disease. Arterioscler Thromb Vasc Biol. 2017;37:218–225.

23. Ionita MG, van den Borne P, Catanzariti LM, Moll FL, de Vries JP, Pasterkamp G, Vink A and de Kleijn DP. High neutrophil numbers in human carotid atherosclerotic plaques are associated with characteristics of rupture-prone lesions. Arterioscler Thromb Vasc Biol. 2010;30:1842–8.

24. Naruko T, Ueda M, Haze K, van der Wal AC, van der Loos CM, Itoh A, Komatsu R, Ikura Y, Ogami M, Shimada Y, Ehara S, Yoshiyama M, Takeuchi K, Yoshikawa J and Becker AE. Neutrophil infiltration of culprit lesions in acute coronary syndromes. Circulation. 2002;106:2894–900.

25. Lacy P. Mechanisms of degranulation in neutrophils. Allergy Asthma Clin Immunol. 2006;2:98–108.

26. Kaiser R, Escaig R, Erber J and Nicolai L. Neutrophil-Platelet Interactions as Novel Treatment Targets in Cardiovascular Disease. Front Cardiovasc Med. 2021;8:824112.

27. Gerhardt T, Haghikia A, Stapmanns P and Leistner DM. Immune Mechanisms of Plaque Instability. Front Cardiovasc Med. 2021;8:797046.

28. Yang LX, Heng XH, Guo RW, Si YK, Qi F and Zhou XB. Atorvastatin inhibits the 5-lipoxygenase pathway and expression of CCL3 to alleviate atherosclerotic lesions in atherosclerotic ApoE knockout mice. J Cardiovasc Pharmacol. 2013;62:205–11.

29. Potteaux S, Combadiere C, Esposito B, Lecureuil C, Ait-Oufella H, Merval R, Ardouin P, Tedgui A and Mallat Z. Role of bone marrow-derived CC-chemokine receptor 5 in the development of atherosclerosis of low-density lipoprotein receptor knockout mice. Arterioscler Thromb Vasc Biol. 2006;26:1858–63.

30. Galkina E, Kadl A, Sanders J, Varughese D, Sarembock IJ and Ley K. Lymphocyte recruitment into the aortic wall before and during development of atherosclerosis is partially L-selectin dependent. J Exp Med. 2006;203:1273–82.

31. Endo-Umeda K, Kim E, Thomas DG, Liu W, Dou H, Yalcinkaya M, Abramowicz S, Xiao T, Antonson P, Gustafsson JA, Makishima M, Reilly MP, Wang N and Tall AR. Myeloid LXR (Liver X Receptor) Deficiency Induces Inflammatory Gene Expression in Foamy Macrophages and Accelerates Atherosclerosis. Arterioscler Thromb Vasc Biol. 2022;42:719–731.

32. Sui Y, Meng Z, Park SH, Lu W, Livelo C, Chen Q, Zhou T and Zhou C. Myeloid-specific deficiency of pregnane X receptor decreases atherosclerosis in LDL receptor-deficient mice. J Lipid Res. 2020;61:696–706.

33. Zhu W, Liang W, Lu H, Chang L, Zhang J, Chen YE and Guo Y. Myeloid TM6SF2 Deficiency Inhibits Atherosclerosis. Cells. 2022;11.

34. Baardman J, Verberk SGS, van der Velden S, Gijbels MJJ, van Roomen C, Sluimer JC, Broos JY, Griffith GR, Prange KHM, van Weeghel M, Lakbir S, Molenaar D, Meinster E, Neele AE, Kooij G, de Vries HE, Lutgens E, Wellen KE, de Winther MPJ and Van den Bossche J. Macrophage ATP citrate lyase deficiency stabilizes atherosclerotic plaques. Nat Commun. 2020;11:6296.

35. Wang F, Liu Z, Park SH, Gwag T, Lu W, Ma M, Sui Y and Zhou C. Myeloid beta-Catenin Deficiency Exacerbates Atherosclerosis in Low-Density Lipoprotein Receptor-Deficient Mice. Arterioscler Thromb Vasc Biol. 2018;38:1468–1478.

36. Sakai K, Nagashima S, Wakabayashi T, Tumenbayar B, Hayakawa H, Hayakawa M, Karasawa T, Ohashi K, Yamazaki H, Takei A, Takei S, Yamamuro D, Takahashi M, Yagyu H, Osuga JI, Takahashi M, Tominaga SI and Ishibashi S. Myeloid HMG-CoA (3-Hydroxy-3-Methylglutaryl-Coenzyme A) Reductase Determines Atherosclerosis by Modulating Migration of Macrophages. Arterioscler Thromb Vasc Biol. 2018;38:2590–2600.

37. Yang G, Zhang J, Jiang T, Monslow J, Tang SY, Todd L, Pure E, Chen L and FitzGerald GA. Bmal1 Deletion in Myeloid Cells Attenuates Atherosclerotic Lesion Development and Restrains Abdominal Aortic Aneurysm Formation in Hyperlipidemic Mice. Arterioscler Thromb Vasc Biol. 2020;40:1523–1532.

38. Gong KQ, Frevert C and Manicone AM. Deletion of LysM in LysMCre Recombinase Homozygous Mice is Non-contributory in LPS-Induced Acute Lung Injury. Lung. 2019;197:819–823.

39. Balta S, Celik T, Mikhailidis DP, Ozturk C, Demirkol S, Aparci M and Iyisoy A. The Relation Between Atherosclerosis and the Neutrophil-Lymphocyte Ratio. Clin Appl Thromb Hemost. 2016;22:405–11.

40. Ruan W, Wang M, Sun C, Yao J, Ma Y, Ma H, Ding J and Lian X. Correlation between neutrophil-to-lymphocyte ratio and stability of carotid plaques. Clin Neurol Neurosurg. 2022;212:107055.

41. Rosales C. Neutrophil: A Cell with Many Roles in Inflammation or Several Cell Types? Front Physiol. 2018;9:113.

42. Kim MH, Yang D, Kim M, Kim SY, Kim D and Kang SJ. A late-lineage murine neutrophil precursor population exhibits dynamic changes during demand-adapted granulopoiesis. Sci Rep. 2017;7:39804.

43. Monaco G, Lee B, Xu W, Mustafah S, Hwang YY, Carre C, Burdin N, Visan L, Ceccarelli M, Poidinger M, Zippelius A, Pedro de Magalhaes J and Larbi A. RNA-Seq Signatures Normalized by mRNA Abundance Allow Absolute Deconvolution of Human Immune Cell Types. Cell Rep. 2019;26:1627–1640 e7.

44. Doring Y, Drechsler M, Wantha S, Kemmerich K, Lievens D, Vijayan S, Gallo RL, Weber C and Soehnlein O. Lack of neutrophil-derived CRAMP reduces atherosclerosis in mice. Circ Res. 2012;110:1052–6.

45. Doring Y, Manthey HD, Drechsler M, Lievens D, Megens RT, Soehnlein O, Busch M, Manca M, Koenen RR, Pelisek J, Daemen MJ, Lutgens E, Zenke M, Binder CJ, Weber C and Zernecke A. Auto-antigenic protein-DNA complexes stimulate plasmacytoid dendritic cells to promote atherosclerosis. Circulation. 2012;125:1673–83.

46. Amersfoort J, Schaftenaar FH, Douna H, van Santbrink PJ, Kroner MJ, van Puijvelde GHM, Quax PHA, Kuiper J and Bot I. Lipocalin-2 contributes to experimental atherosclerosis in a stage-dependent manner. Atherosclerosis. 2018;275:214–224.

47. Paclet MH, Laurans S and Dupre-Crochet S. Regulation of Neutrophil NADPH Oxidase, NOX2: A Crucial Effector in Neutrophil Phenotype and Function. Front Cell Dev Biol. 2022;10:945749.

48. Kim K, Li J, Tseng A, Andrews RK and Cho J. NOX2 is critical for heterotypic neutrophil-platelet interactions during vascular inflammation. Blood. 2015;126:1952–64.

49. Nijm J, Wikby A, Tompa A, Olsson AG and Jonasson L. Circulating levels of proinflammatory cytokines and neutrophil-platelet aggregates in patients with coronary artery disease. Am J Cardiol. 2005;95:452–6.

50. Zhou J, Xu E, Shao K, Shen W, Gu Y, Li M and Shen W. Circulating platelet-neutrophil aggregates as risk factor for deep venous thrombosis. Clin Chem Lab Med. 2019;57:707–715.

51. Pircher J, Czermak T, Ehrlich A, Eberle C, Gaitzsch E, Margraf A, Grommes J, Saha P, Titova A, Ishikawa-Ankerhold H, Stark K, Petzold T, Stocker T, Weckbach LT, Novotny J, Sperandio M, Nieswandt B, Smith A, Mannell H, Walzog B, Horst D, Soehnlein O, Massberg S and Schulz C. Cathelicidins prime platelets to mediate arterial thrombosis and tissue inflammation. Nat Commun. 2018;9:1523.

52. Filippi MD. Neutrophil transendothelial migration: updates and new perspectives. Blood. 2019;133:2149–2158.

53. Drechsler M, Megens RT, van Zandvoort M, Weber C and Soehnlein O. Hyperlipidemia-triggered neutrophilia promotes early atherosclerosis. Circulation. 2010;122:1837–45.

54. Hartl D, Krauss-Etschmann S, Koller B, Hordijk PL, Kuijpers TW, Hoffmann F, Hector A, Eber E, Marcos V, Bittmann I, Eickelberg O, Griese M and Roos D. Infiltrated neutrophils acquire novel chemokine receptor expression and chemokine responsiveness in chronic inflammatory lung diseases. J Immunol. 2008;181:8053–67.

55. de Jager SC, Bot I, Kraaijeveld AO, Korporaal SJ, Bot M, van Santbrink PJ, van Berkel TJ, Kuiper J and Biessen EA. Leukocyte-specific CCL3 deficiency inhibits atherosclerotic lesion development by affecting neutrophil accumulation. Arterioscler Thromb Vasc Biol. 2013;33:e75–83.

56. Mawhin MA, Tilly P, Zirka G, Charles AL, Slimani F, Vonesch JL, Michel JB, Back M, Norel X and Fabre JE. Neutrophils recruited by leukotriene B4 induce features of plaque destabilization during endotoxaemia. Cardiovasc Res. 2018;114:1656–1666.

57. Burn GL, Foti A, Marsman G, Patel DF and Zychlinsky A. The Neutrophil. Immunity. 2021;54:1377–1391.

